# *trans*-Translation inhibitors that kill *M. tuberculosis* and pathogenic Non-tuberculous Mycobacteria have a dual mechanism of action

**DOI:** 10.64898/2026.01.22.701086

**Authors:** Akanksha Varshney, Ziyi Jia, Gebremichal Gebretsadik, Narendran G-Dayanandan, Terry L. Bowlin, Michelle M. Butler, Anthony D. Baughn, Kenneth C. Keiler

## Abstract

*Mycobacterium tuberculosis* and pathogenic nontuberculous mycobacteria pose a growing challenge to human health, and new antibiotics that target new pathways with novel mechanisms of action are urgently needed. Acylaminooxadiazole derivatives have previously been shown to inhibit the *trans-*translation ribosome rescue pathway and kill *M. tuberculosis*. Here, we show that modifications to the acylaminooxadiazole scaffold can improve potency and tune mycobacterial species specificity, resulting in molecules that kill *M. avium*, *M. abscessus*, and *M. tuberculosis* clinical isolates. Free iron was previously shown to antagonize antibacterial activity and decrease the inhibition of *trans*-translation by acylaminooxadiazoles, but we found that biologically relevant iron sources such as hemin and transferrin do not affect activity. Mutants depleted for tmRNA and mutants defective in siderophore-mediated iron utilization are both hypersusceptible to acylaminooxadiazole-based *trans-*translation inhibitors, indicating a dual mechanism of action involving both direct inhibition of *trans-*translation and metal starvation. These findings establish acylaminooxadiazoles as dual-mechanism antimycobacterial agents that couple inhibition of *trans*-translation with disruption of iron homeostasis.

## INTRODUCTION

*Mycobacterium tuberculosis*, the etiological agent of tuberculosis (TB), remains a leading cause of infectious disease-related mortality worldwide. In 2024, an estimated 10.6 million new TB cases and 1.3 million deaths were reported, reflecting the continued global burden of this pathogen [1]. Although recently introduced multidrug regimens and shorter treatment courses have shown encouraging success against drug-susceptible and some drug-resistant forms of TB, the emergence and spread of multidrug-resistant (MDR) and extensively drug-resistant (XDR) strains continue to threaten these gains [2–4]. In parallel, non-tuberculous mycobacteria (NTM), such as *Mycobacterium avium* and *Mycobacterium abscessus*, have emerged as important opportunistic pathogens that cause chronic pulmonary and disseminated infections, particularly in immunocompromised patients [5, 6]. NTM infections are intrinsically drug resistant and require prolonged multidrug therapy, highlighting the need for new antimycobacterial agents with novel mechanisms of action effective against both *M. tuberculosis* and NTM [7–11].

Essential bacterial processes that are absent in humans are especially attractive targets for new antimycobacterial drugs because they can offer both efficacy and selectivity. One such process is *trans*-translation, a ribosome rescue system that resolves “non-stop” translation complexes in bacteria [12, 13]. Non-stop complexes arise when ribosomes reach the 3′ end of an mRNA without terminating at a stop codon and become stalled [12, 13]. In these events, the tmRNA–SmpB complex, which consists of transfer-messenger RNA (tmRNA) and small protein B (SmpB), enters the vacant A site of the stalled ribosome. This complex mimics both tRNA and mRNA to initiate rescue [14–16]. tmRNA first participates in a transpeptidation reaction in which the nascent peptide is transferred to an alanine attached to tmRNA. It then provides a short internal open reading frame encoding a degradation tag [12, 13]. Translation resumes on this tmRNA template, adding the tag to the nascent chain and terminating normally [12, 13]. The tagged protein is subsequently recognized and degraded by cellular proteases, while the ribosome is recycled for new rounds of translation [12, 17–19]. This quality-control system is essential in *M. tuberculosis* and is absent from human cells, making it an especially compelling antibacterial target [20, 21].

We have identified several classes of small-molecule inhibitors of *trans*-translation that kill *M. tuberculosis* through distinct mechanisms. Acylaminooxadiazoles, such as KKL-35 and MBX-4132, act by targeting the 23S rRNA [22, 21, 23, 24]. Tetrazole derivatives, including KKL-55, inhibit *trans*-translation through binding to elongation factor Tu, while triazole analogs, such as KKL-1005, act by targeting ribosomal protein L12 [25, 26]. Intriguingly, RNA-sequencing and transposon insertion sequencing analyses of *M. tuberculosis* treated with the 1,3,4-acylaminooxadiazole, MBX-4132, showed that the cells respond as though they were starved for iron and experienced broad metal dysregulation. These phenotypes suggest the compounds not only inhibit *trans*-translation but also disrupt bacterial metal homeostasis. Additionally, when these compounds were tested in culture, zinc supplementation enhanced antibacterial activity, whereas free iron antagonized this activity [24]. Building on these findings, we tested whether biologically relevant forms of iron influence compound activity and whether acylaminooxadiazole based *trans*-translation inhibitors have a dual mechanism of action.

In this study, we developed new acylaminooxadiazole analogs with improved antimycobacterial potency and enhanced on-target inhibition of *trans*-translation in *M. tuberculosis* and clinically relevant NTM species. We also assessed how physiologically relevant forms of iron affect acylaminooxadiazole-mediated inhibition of *trans*-translation. Here, we describe the biochemical and microbiological characterization of analogs derived from the KKL-35 and MBX-4132 scaffolds. Collectively, our findings expand the acylaminooxadiazole class and highlight distinct structure–activity relationships that govern both inhibition of *trans*-translation and modulation of metal homeostasis. These insights underscore the biological relevance of metal interactions in modulating inhibitor activity and provide a framework for optimizing this scaffold for therapeutic development.

## RESULTS

### Species specific antimycobacterial activity by acylaminooxadiazoles

We evaluated a series of 1,3,4- and 1,2,4-acylaminooxadiazole analogs based on the MBX-4132 scaffold to identify compounds with improved potency against *M. tuberculosis*, *M. avium*, and *M. abscessus* (Figure 1). Structural modifications were introduced at both the oxadiazole core and the adjacent rings to examine how these changes affect efficacy and inhibition of *trans*-translation. Because previous work showed that excess iron reduced acylaminooxadiazole activity in liquid culture, we performed MIC and MBC assays in low-iron Mtb Minimal Medium (LIMM) to minimize iron interference and accurately assess intrinsic potency [24]. Several compounds exhibited greater antimycobacterial activity than MBX-4132. MBX-5822A (MBC = 0.2 µg/mL) was the most potent against *M. tuberculosis* H37Rv Δ*RD1* Δ*panCD*, followed by MBX-6958A (MBC = 0.4 µg/mL), both outperforming MBX-4132 in MBC assays (Table 1). For comparison, first-line drugs such as rifampicin (RIF) and isoniazid (INH) have MBCs in a similar range (≈0.125–0.5 µg/mL), so MBX-5822A and MBX-6958A fall within the potency range of clinically effective antibiotics (Table 1) [27].

**Figure 1.**
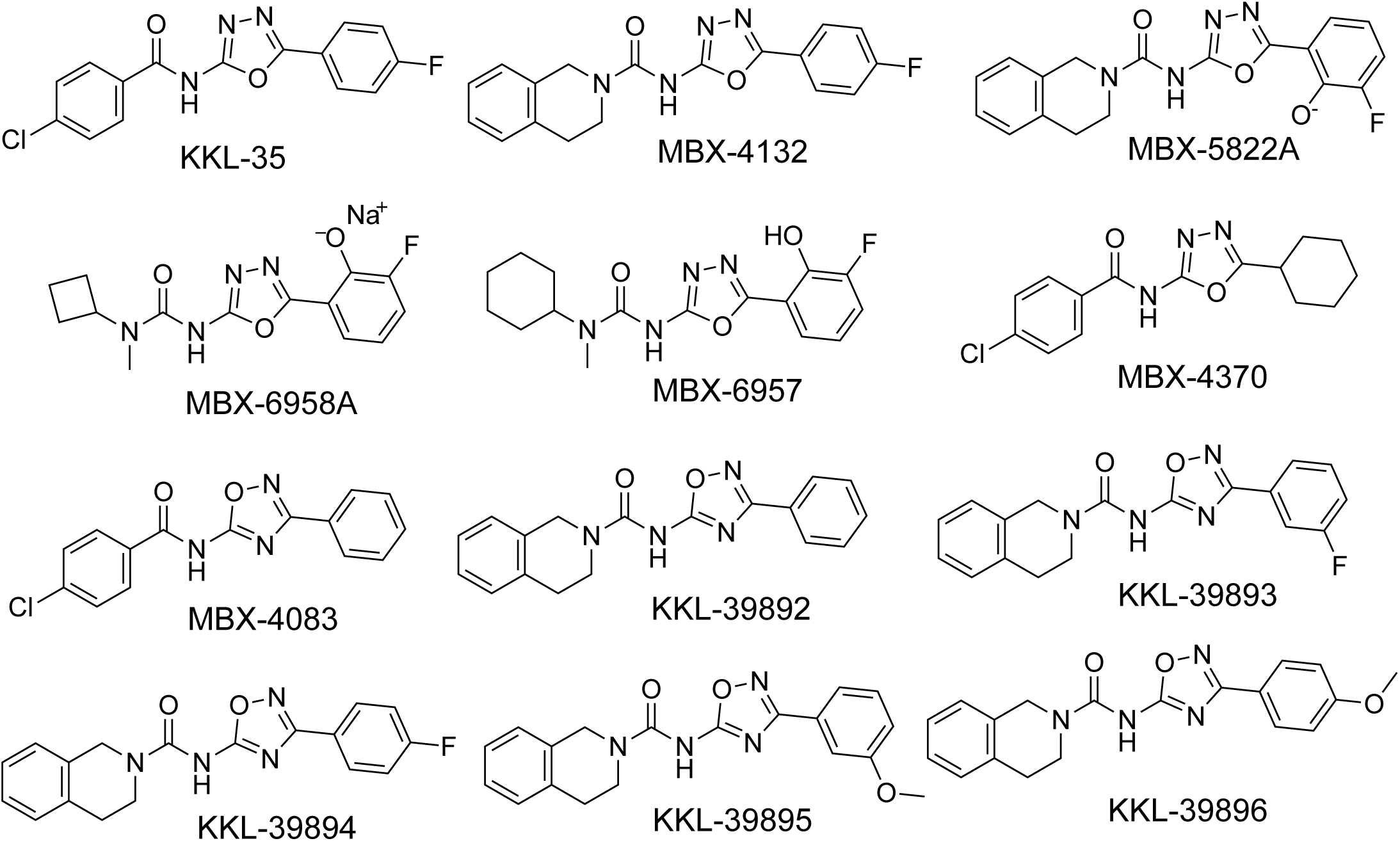
Structures of acylaminooxadiazoles used in this study.

**Table 1.** Minimum inhibitory and minimum bactericidal concentrations of *trans-*translation inhibitors for mycobacterial species in LIMM.

| Compound | *M. tuberculosis*<br>H37Rv $\Delta RD1$<br>$\Delta panCD$ | | *M. avium* | | *M. abscessus* | |
| --- | --- | --- | --- | --- | --- | --- |
|  | MIC<sup>a</sup> | MBC<sup>b</sup> | MIC | MBC | MIC | MBC |
| KKL-35 | 0.4<br>(1.25) | 0.4<br>(1.25) | 3.1<br>(9.7) | 3.1<br>(9.7) | 6.3<br>(19.8) | 6.3<br>(19.8) |
| MBX-4132 | 0.8<br>(2.5) | 1.6<br>(5) | 2.1<br>(6.2) | 2.1<br>(6.2) | 3.3<br>(9.7) | 3.3<br>(9.7) |
| MBX-5822A | 0.23<br>(0.62) | 0.23<br>(0.62) | 3.75<br>(10) | 3.75<br>(10) | 7.5<br>(20) | 7.5<br>(20) |
| MBX-6958A | 0.41<br>(1.25) | 0.23<br>(0.62) | 4.65<br>(10) | 4.65<br>(10) | 9.3<br>(20) | 9.3<br>(20) |
| MBX-6957 | 0.23<br>(1.25) | 0.23<br>(1.25) | 3.75<br>(0.31) | 3.75<br>(0.31) | 1.67<br>(5) | 1.67<br>(5) |
| MBX-4370 | 0.30<br>(1.25) | 0.3<br>(0.62) | 0.12<br>(0.41) | 0.12<br>(0.41) | 1.52<br>(5) | 1.52<br>(5) |
| MBX-4083 | 0.8<br>(25) | 0.8<br>(25) | 24<br>(75) | 24<br>(75) | 64<br>(200) | 64<br>(200) |
| KKL-39892 | 32<br>(100) | 16<br>(50) | >32<br>(>100) | >32<br>(>100) | >32<br>(>100) | >32<br>(>100) |
| KKL-39893 | 4<br>(12.5) | 4<br>(12.5) | 32<br>(100) | 32<br>(100) | >32<br>(>100) | >32<br>(>100) |
| KKL-39894 | 2<br>(6.25) | 2<br>(6.25) | 16<br>(50) | 16<br>(50) | >32<br>(>100) | >32<br>(>100) |
| KKL-39895 | 16<br>(50) | 16<br>(50) | >32<br>(>100) | >32<br>(>100) | >32<br>(>100) | >32<br>(>100) |
| KKL-39896 | 4<br>(12.5) | 4<br>(12.5) | 32<br>(100) | 32<br>(100) | >32<br>(>100) | >32<br>(>100) |
| Rifampicin | 0.12<br>(0.14) | 0.12<br>(0.14) | 0.5<br>(0.6) | 1<br>(1.2) | 128<br>(155) | 256<br>(311) |
| Azithromycin | 4<br>(5.3) | ND<sup>c</sup> | 32<br>(42.4) | ND | 8<br>(10.6) | ND |

<sup>a</sup>μg/mL (μM) values from at least three broth microdilution assays.
<sup>b</sup>μg/mL (μM) values from at least three plating assays.
<sup>c</sup>Not determined.

*M. avium* is intrinsically resistant to most current clinical antibiotics, and only a small number of effective agents achieve MBCs of ≤4 µg/mL [28, 29], MBX-5822A (MBC = 3.7 µg/mL) and MBX-6958A (MBC = 4.6 µg/mL) demonstrated more potent activity than MBX-4132 (MBC = 9.0 µg/mL). Notably, MBX-4370 (MBC = 0.1 µg/mL) was the most active compound against this species, indicating that modifications to the aromatic and amide side chains while retaining the core 1,3,4-oxadiazole scaffold, can markedly enhance activity against NTM (Table 1).

*M. abscessus* is among the most drug-resistant nontuberculous mycobacteria, and compounds with MBCs ≤8 µg/mL are generally considered highly active against this pathogen [11, 30]. MBX-5822A (MBC = 7.5 µg/mL) and MBX-6958A (MBC = 9.3 µg/mL) showed more potent activity than MBX-4132 against *M. abscessus*, whereas MBX-4370 (MBC = 1.5 µg/mL) exhibited exceptional potency, falling within the range reported for the most effective antibacterial agents against this pathogen (Table 1).

In contrast to the 1,3,4-acylaminooxadiazoles, all the tested 1,2,4-acylaminooxadiazoles exhibited lower potency than MBX-4132. Within this class, MBX-4083 displayed the highest potency and was bactericidal against *M. tuberculosis* H37Rv Δ*RD1* Δ*panCD* (MBC = 4 µg/mL), *M. avium* (MBC = 8 µg/mL), and *M. abscessus* (MBC = 15 µg/mL) (Table 1). These results confirm that the 1,3,4-oxadiazole conformation is important for *trans*-translation inhibition, consistent with previous data [23].

Because clinical isolates of the *M. tuberculosis* complex (MTBC) often exhibit variable drug susceptibility, we assessed whether MBX-4132 and MBX-5822A retained activity against a panel of these strains. 10 such strains were included, representing 6 out of the 9 MTBC phylogenetic lineages, in addition to H37Rv for reference [31]. Compound efficacy was evaluated in both Middlebrook 7H9 (containing ∼6 µM ZnSO_4_) and high zinc 7H9, prepared as 7H9 supplemented with 100 µM ZnSO_4_. We observed that most clinical isolates displayed similar or slightly enhanced susceptibility to the compounds compared to H37Rv under both conditions (Table 2, Figure S1). We also found that N1274 was the most susceptible strain to MBX-4132 and isoniazid (INH, control) in both media and MBX-5822A only in standard 7H9 (Table 2, Figure S1). N1274 is a lineage 3 clinical isolate that is resistant to streptomycin, so its increased susceptibility to these oxadiazoles underscores that activity is maintained even in some strains carrying pre-existing antibiotic resistance [31]. These results support the potential of MBX-4132 and MBX-5822A against diverse MTBC infections, including drug resistant ones. Overall, these results show that changes to MBX-4132 can improve efficacy against *M. tuberculosis* and NTM, and that the compounds with highest activity against each species were distinct.

**Table 2.** Minimum inhibitory concentrations of acylaminooxadiazoles against MTBC clinical isolates.

| Strain | Lineage<sup>a</sup> | DR<sup>b</sup> | MIC in 7H9<sup>c</sup> |  |  | MIC in 7H9+100 µM ZnSO<sub>4</sub><sup>c</sup> |  |  |
| --- | --- | --- | --- | --- | --- | --- | --- | --- |
|  |  |  | INH | MBX-4132 | MBX-5822A | INH | MBX-4132 | MBX-5822A |
| H37Rv | L4 | WT | 0.34<br>(2.5) | 15<br>(50) | 38<br>(100) | 0.34<br>(2.5) | 8<br>(25) | 19<br>(50) |
| N0004 | L3 | WT | 1.36<br>(10) | 17<br>(50) | 38<br>(100) | 0.68<br>(5) | 17<br>(50) | 19<br>(50) |
| N0052 | L2 | WT | >0.14<br>(>1) | 34<br>(100) | 38<br>(100) | 0.34<br>(2.5) | 17<br>(50) | 38<br>(100) |
| N0136 | L4 | WT | >0.14<br>(>1) | 8<br>(25) | 38<br>(100) | >0.14<br>(>1) | 4<br>(12.5) | 19<br>(50) |
| N0145 | L2 | WT | 0.07<br>(0.5) | 17<br>(50) | 38<br>(100) | 0.07<br>(0.5) | 4<br>(12.5) | 9<br>(25) |
| N0155 | L2 | WT | 0.34<br>(2.5) | 34<br>(100) | 38<br>(100) | 0.17<br>(1.25) | 4<br>(12.5) | 9<br>(25) |
| N1063 | L5 | WT | >0.14<br>(>1) | >34<br>(>100) | >38<br>(>100) | >0.14<br>(>1) | 17<br>(50) | 19<br>(50) |
| N1202 | L6 | WT | 0.34<br>(2.5) | 34<br>(100) | 38<br>(100) | 0.08<br>(0.63) | 8<br>(25) | 5<br>(12.5) |
| N1216 | L4 | WT | 0.34<br>(2.5) | 34<br>(100) | 38<br>(100) | 0.17<br>(1.25) | 4<br>(12.5) | 5<br>(12.5) |
| N1272 | L5 | WT | 0.07<br>(0.5) | 34<br>(100) | 38<br>(100) | 0.07<br>(0.5) | 17<br>(50) | 38<br>(100) |
| N1274 | L3 | STR | 0.07<br>(0.5) | 8<br>(25) | 9<br>(25) | 0.03<br>(0.25) | 2<br>(6.25) | 9<br>(25) |

<sup>a</sup>Lineage assignments of clinical isolates as described by Borrell et al. and Stucki et al. [31, 40]
<sup>b</sup>Phenotypic drug resistance (DR) profile, indicating either wild-type (WT) or streptomycin (STR)-resistant phenotype based on Borrell et al. and Guerrero-
Bustamante et al. [31, 41]
<sup>c</sup>µg/mL (µM) values from two broth microdilution assays.

### Acylaminooxadiazoles inhibit *M. tuberculosis trans*-translation cell-free and whole cell assays

Since MBX-5822A and MBX-6958A showed greater bactericidal activity than MBX-4132, we next examined whether their inhibition of *trans*-translation was also enhanced. We used our previously established *M. tuberculosis* cell-free *trans*-translation assay containing purified ribosomes, tmRNA, SmpB, and translation factors from *M. tuberculosis* [24, 26]. As previously observed with this assay, active *trans*-translation in the absence of an inhibitor produced both the full-length DHFR protein (DHFR-ns) and a higher-molecular-weight DHFR species containing the tmRNA-encoded peptide tag (Figure 2) [24, 26]. The higher molecular weight band was shown to be the result of tagging as the abundance of this band decreases substantially when tmRNA–SmpB is omitted from the reaction [24, 26]. Addition of MBX-5822A or MBX-6958A reduced the amount of tagged DHFR in a dose-dependent manner, with IC₅₀ values of 2 ± 1 µM and 3 ± 0.6 µM, respectively (Figure 2A-D). MBX-4083 also inhibited *trans*-translation, though less potently, with an IC₅₀ of 21 ± 2 µM (Figure 2E &F). To assess the effects of MBX-5822A, MBX-6958A, and MBX-4083 on normal translation, we employed the *in vitro* translation reactions as described previously [24]. The reaction mixture was programmed with a DHFR template containing an in-frame stop codon (*dhfr-stop*), and tmRNA–SmpB were omitted from the reaction. Under these conditions, none of the tested acylaminooxadiazoles inhibited translation of DHFR-stop at concentrations up to 90 µM, whereas chloramphenicol completely blocked the synthesis of DHFR-stop (Figure 2G). These results show that MBX-5822A, MBX-6958A, and MBX-4083 inhibit *M. tuberculosis trans*-translation and not canonical translation.

**Figure 2.**
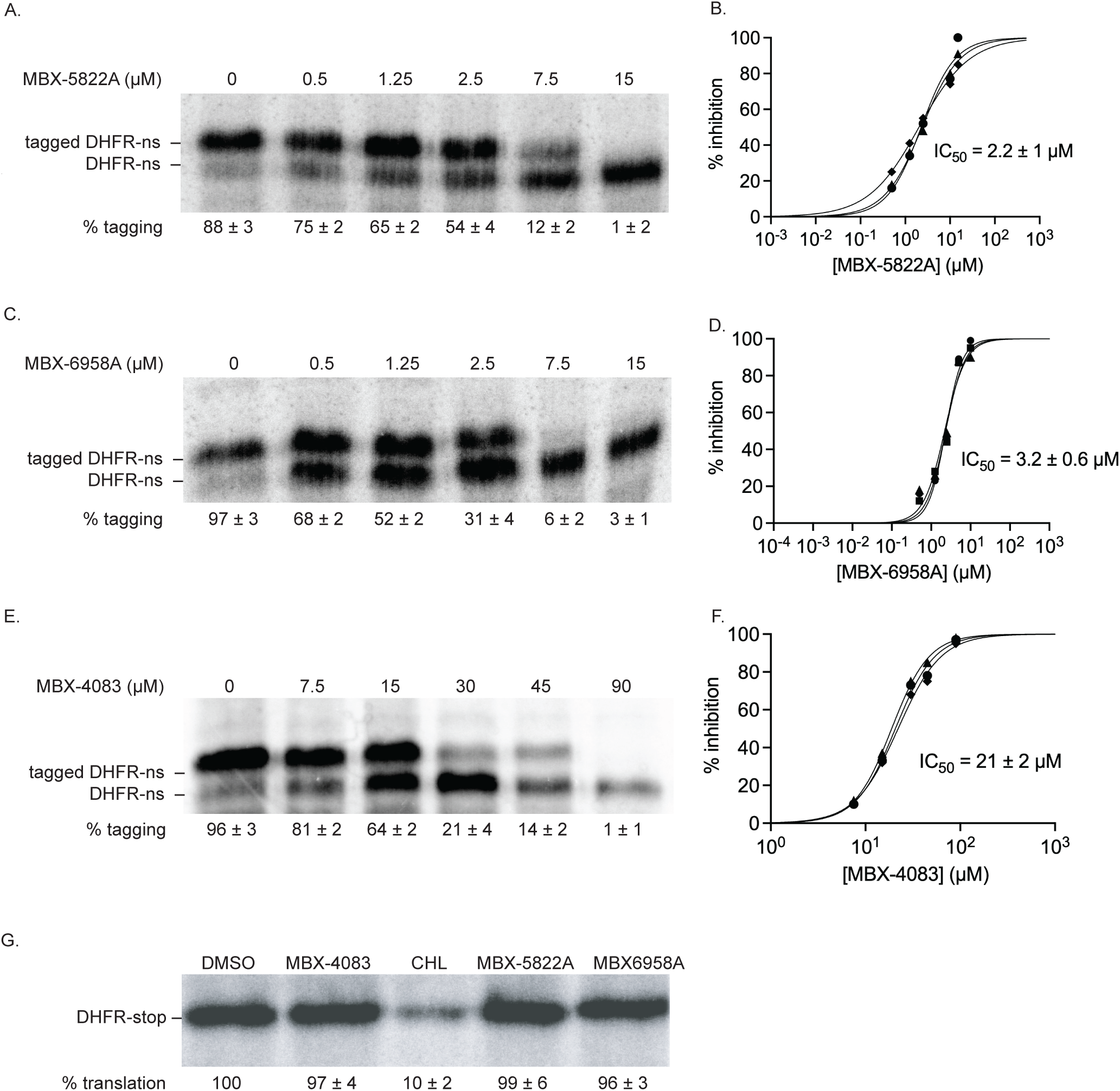
Acylaminooxadiazoles inhibit *M. tuberculosis trans*-translation *in vitro*. A) A gene encoding DHFR without a stop codon (DHFR-ns) was expressed in the presence of *M*. *tuberculosis* tmRNA-SmpB and varying concentrations of MBX-5822A. Synthesized protein was detected by incorporation of ^35^S-methionine followed by SDS-PAGE and phosphorimaging. Bands corresponding to DHFR-ns with tagged DHFR, which is the product of trans-translation are indicated, and the average percentage of DHFR protein found in the tagged band for three repeats is shown with the standard deviation. B) Data from gels as in (A) were plotted and fit with a sigmoidal function to determine the IC_50_. C) *in vitro trans*-translation reactions as in (A) containing varying concentrations of MBX-6958A. D) Data from gels as in (C) were plotted and fit with a sigmoidal function to determine the IC_50._ E*) in vitro* trans-translation reactions as in (A) containing varying concentrations of MBX-4083. F) Data from gels as in (E) were plotted and fit with a sigmoidal function to determine the IC_50._ G) *in vitro* translation was assayed from the expression of a gene encoding DHFR with a stop codon (DHFR-stop) in the presence of DMSO, 20 μM chloramphenicol (CHL), or 20 μM MBX-5822A or MBX-6958A or MBX-4083, and a representative experiment is shown. The percentage of DHFR with respect to the amount in the DMSO-treated control is shown as the average from two independent repeats with the standard deviation.

We next tested whether MBX-5822A and MBX-6958A compete with tmRNA–SmpB for binding to the ribosome, as previously observed for MBX-4132 [24]. Using the *M. tuberculosis* cell-free *trans*-translation assay in the presence of increasing concentrations of tmRNA–SmpB, we found that higher tmRNA–SmpB concentrations reduced inhibition of *trans*-translation and increasing the compound concentration restored inhibition (Figure 3A-D). These data indicate that MBX-5822A and MBX-6958A compete with tmRNA-SmpB activity on the *M. tuberculosis* ribosomes, and that the structural changes to MBX-4132 did not alter inhibition of *trans*-translation (Figure 3A-D).

**Figure 3.**
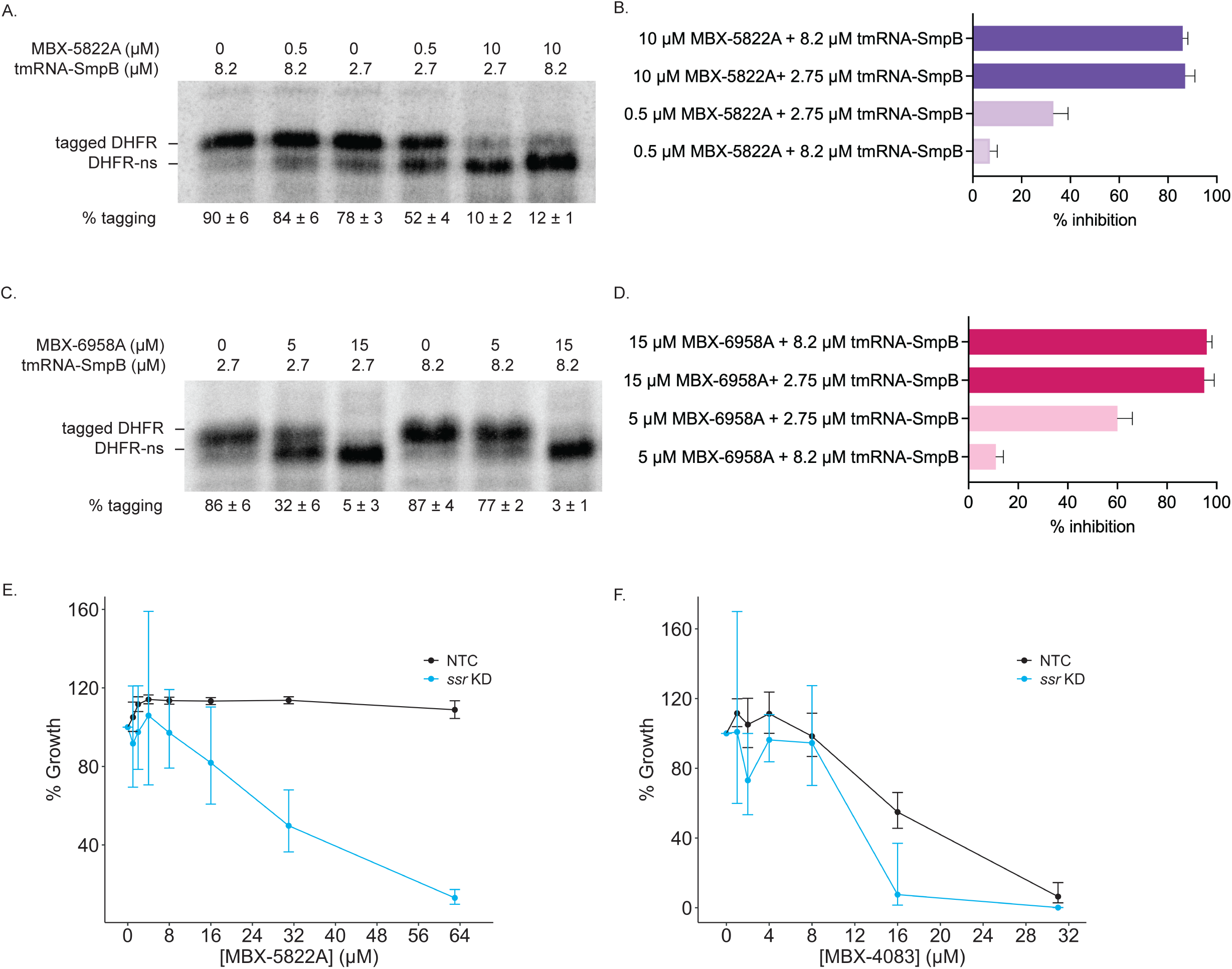
tmRNA-SmpB compete with acylminooxadiazoles *in vitro.* A) *in vitro trans*-translation assays as in Figure 2 containing different concentrations of tmRNA-SmpB and MBX-5822A. B) Reactions treated with 8.2 μM tmRNA-SmpB suppressed the inhibition of *trans*-translation by 0.5 μM MBX-4132. The inhibition was rescued by 10 μM MBX-5822A. Data from at least two experiments are shown as the average with error bars indicating the standard deviation. C) *in vitro trans*-translation reactions as in (A) containing different concentrations of tmRNA-SmpB and MBX-5822A. D) Reactions treated with 8.2 μM tmRNA-SmpB suppressed the inhibition of *trans*-translation by 5 μM MBX-6958A. The inhibition was rescued by 15 μM MBX-6958A. E-F) Susceptibility of *M. tuberculosis* H37Rv *ssr* KD (blue) and NTC (black) strains to (E) MBX-5822A and (F) MBX-4083 Cells were treated with ATc for 7 days and subsequently exposed to acylaminooxadiazoles for 21 days in MM. Data represent geometric means and geometric standard deviations for 3 biological replicates.

Our previous research established that CRISPRi-mediated knockdown of *ssr* increased the sensitivity of *M. tuberculosis* H37Rv to MBX-4132 compared to the non-targeting control strain [24, 32]. Here, we tested whether reduced tmRNA levels had a similar effect on MBX-5822A and MBX-4083. We observed that the hypomorph conferred enhanced susceptibility to both compounds by several-fold, albeit to a lesser extent than MBX-4132 (Figure 3E & F).

### Acylaminooxadiazole based *trans*-translation inhibitors are not antagonistic to existing antimycobacterial antibiotics

Since new compounds may be used alongside existing antimycobacterial therapies, we evaluated whether acylaminooxadiazoles exhibit antagonistic interactions with clinically used antimycobacterial drugs. For example, RIF is known to antagonize the activity of certain β-lactams against *M. tuberculosis* by inducing drug-metabolizing enzymes and altering bacterial physiology [33]. To evaluate whether the acylaminooxadiazoles exhibit antagonism, we tested MBX-4132, MBX-5822A, and MBX-6958A in combination with some of the clinically relevant antitubercular agents against *M. tuberculosis* H37Rv Δ*RD1* Δ*panCD* using checkerboard MIC assays (Table 3). Fractional inhibitory concentration indices (FICI) were determined for each combination, and values were interpreted as synergistic (FICI ≤ 0.5), additive (0.5 < FICI ≤ 2.0), indifferent (2.0 < FICI ≤ 4.0), or antagonistic (FICI > 4.0). MBX-4132, MBX-5822A, and MBX-6958A displayed additive interactions across all combinations (0.5 < FICI ≤ 1.0), consistent with an independent or complementary mechanism of action (Table 3). We next evaluated MBX-4132, MBX-5822A, and MBX-6958A for potential antagonism with some of the clinically used agents against *M. avium*, including azithromycin, erythromycin, and linezolid (LNZ). None of the compounds showed antagonistic interactions with any of these tested drugs (Table 4). Together, these data indicate that MBX-4132, MBX-5822A, and MBX-6958A are not antagonistic to commonly used antimycobacterial drugs and may be compatible with existing multidrug treatment regimens for TB and NTM infections.

**Table 3.**
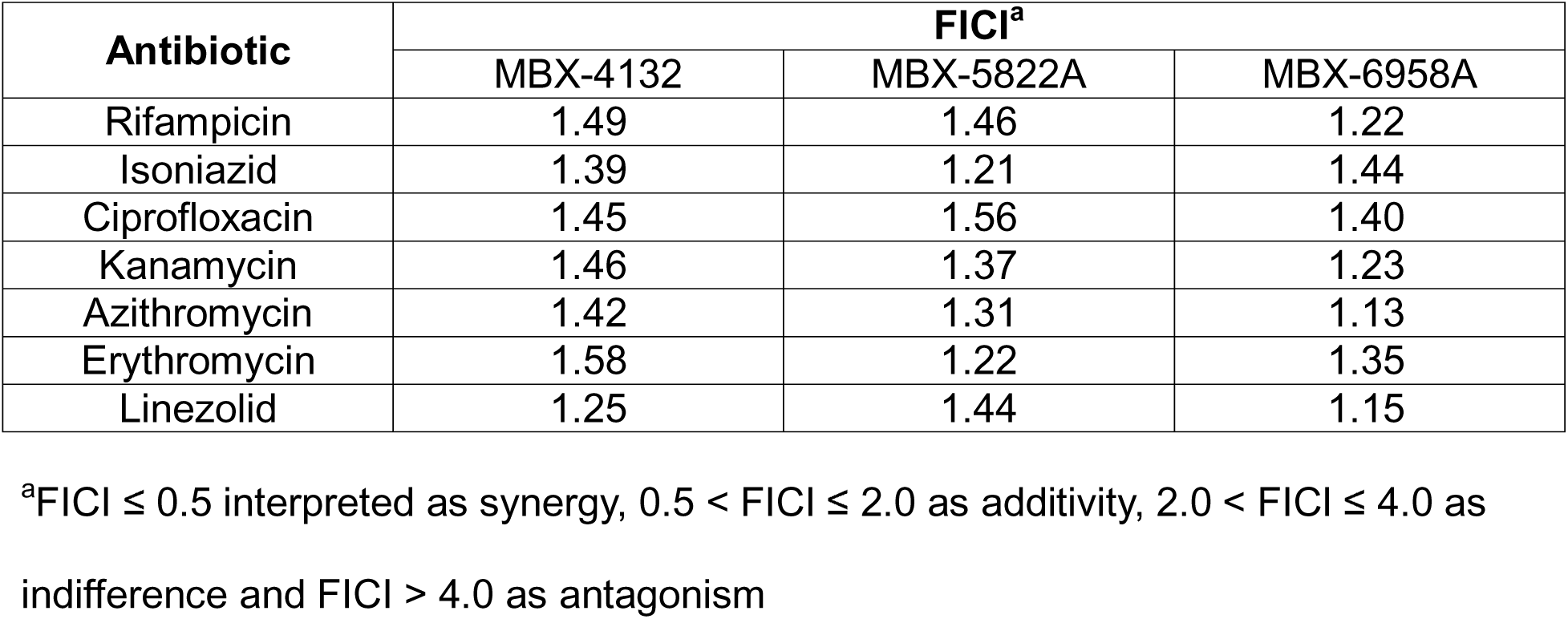
Checkerboard MIC assay to evaluate drug-combinations against *M. tuberculosis* H37Rv Δ*RD1* Δ*panCD*.

| Antibiotic | FICI<sup>a</sup> |  |  |
| --- | --- | --- | --- |
|  | MBX-4132 | MBX-5822A | MBX-6958A |
| Rifampicin | 1.49 | 1.46 | 1.22 |
| Isoniazid | 1.39 | 1.21 | 1.44 |
| Ciprofloxacin | 1.45 | 1.56 | 1.40 |
| Kanamycin | 1.46 | 1.37 | 1.23 |
| Azithromycin | 1.42 | 1.31 | 1.13 |
| Erythromycin | 1.58 | 1.22 | 1.35 |
| Linezolid | 1.25 | 1.44 | 1.15 |

<sup>a</sup>FICI $\leq 0.5$ interpreted as synergy, $0.5 < \text{FICI} \leq 2.0$ as additivity, $2.0 < \text{FICI} \leq 4.0$ as indifference and $\text{FICI} > 4.0$ as antagonism

**Table 4.** Checkerboard MIC assays to evaluate drug-combinations against *M. avium*.

| Antibiotic | FICI<sup>a</sup> |  |  |
| --- | --- | --- | --- |
|  | MBX-4132 | MBX-5822A | MBX-6958A |
| Azithromycin | 1.36 | 1.29 | 1.30 |
| Erythromycin | 1.46 | 1.26 | 1.21 |
| Linezolid | 1.12 | 1.22 | 1.14 |

<sup>a</sup>FICI $\leq 0.5$ interpreted as synergy, $0.5 < \text{FICI} \leq 2.0$ as additivity, $2.0 < \text{FICI} \leq 4.0$ as indifference and $\text{FICI} > 4.0$ as antagonism

### Biologically complexed iron does not antagonize the antimycobacterial activity of acylaminooxadiazoles

We previously found that free iron impairs the activity of MBX-4132 in cell-free and culture-based assays, suggesting that the metal directly binds to MBX-4132 and prevents it from interacting with the ribosome [24]. To determine if free iron also interferes with MBX-5822A, MBX-6958A, and MBX-4083, we repeated the *in vitro trans*-translation and growth inhibition assays in the presence of free iron. Similar to MBX-4132, free iron prevented MBX-5822A and MBX-6958A from inhibiting *trans*-translation *in vitro* (Figure 4). Likewise, MBX-5822A and MBX-6958A did not inhibit the growth of *M. tuberculosis* or NTMs in 7H9 medium that contained free iron (Table S1). In contrast, MBX-4083 activity was not affected by free iron *in vitro* or in culture (Figure 4, Table S1).

**Figure 4.**
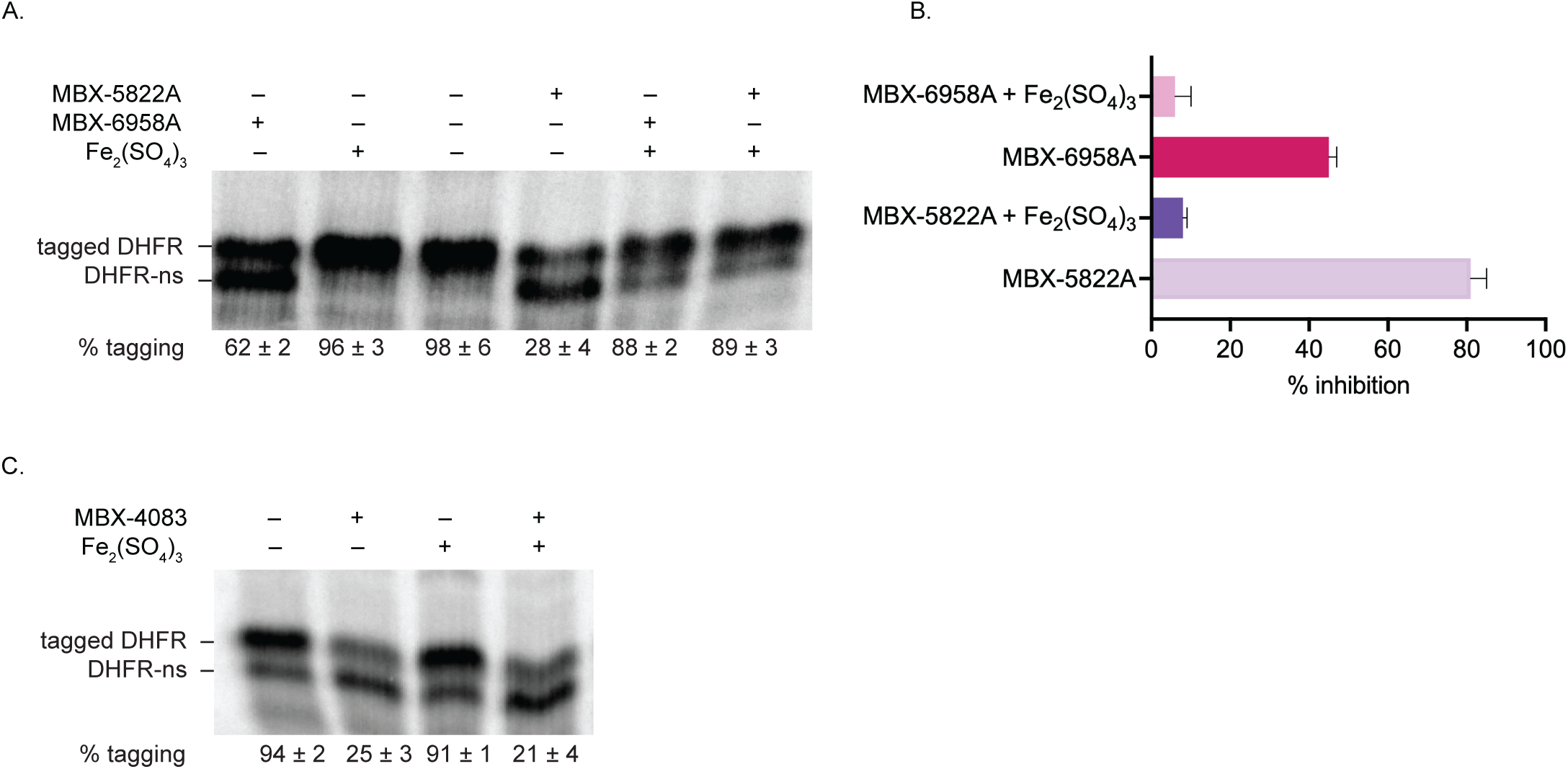
Free iron antagonizes the activity of MBX-5822A and MBX-6958A but not MBX-4083. A) *in vitro trans*-translation assays as in Figure 2 containing 150 μM Fe_2_(SO_4_)_3_, 15 μM MBX-5822A, or MBX-6958A, as indicated. The average percentage tagging from two independent reactions is shown with the standard deviation. B) Data from gels as in (A) were plotted to show the average from two experiments with error bars indicating the standard deviation. C) *in vitro trans-*translation assays as in Figure 2 containing 200 μM Fe_2_(SO_4_)_3_, 20 μM MBX-4083, as indicated.

To assess whether other transition metals also influence oxadiazole activity, we examined the effects of copper and zinc on activity of acylaminooxadiazoles. Copper supplementation increased the MICs of MBX-4132, KKL-35, MBX-5822A, and MBX-6958A in *M. tuberculosis* Δ*RD1* Δ*panCD* cultures, indicating reduced antibacterial activity (Table S2). Consistent with this finding, copper also decreased inhibition of *trans*-translation by MBX-4132 and KKL-35 *in vitro*, and addition of the metal chelator TPEN restored activity, confirming that Cu²⁺ directly interferes with compound function (Figure 5). Together, these data demonstrate that copper antagonizes acylaminooxadiazole activity both in culture and biochemically. In contrast, increased zinc potentiated the activity of 1,3,4-and 1,2,4-acylaminooxadiazoles in liquid cultures, as previously observed for MBX-4132 [24]. MBX-5822A and MBX-6958A had substantially lower MICs in high-zinc minimal medium (HZMM) compared to MM, whereas MBX-4083 activity was unchanged (Table S3). When copper was added to HZMM, the activity of MBX-4132, MBX-5822A, and MBX-6958A was impaired, but the activity of MBX-4083 was not affected. Taken together, these data suggest that the 1,3,4-acylaminooxadizoles bind free iron and copper and cannot inhibit *trans-*translation in the metal-bound state, whereas MBX-4083 does not bind iron or copper. Potentiation by zinc is likely to be the result of a physiological response by the bacteria because zinc does not have any effect *in vitro.* The basis for how structural differences among these acylaminooxadiazoles influence metal modulation remains unclear and will require further investigation.

**Figure 5.**
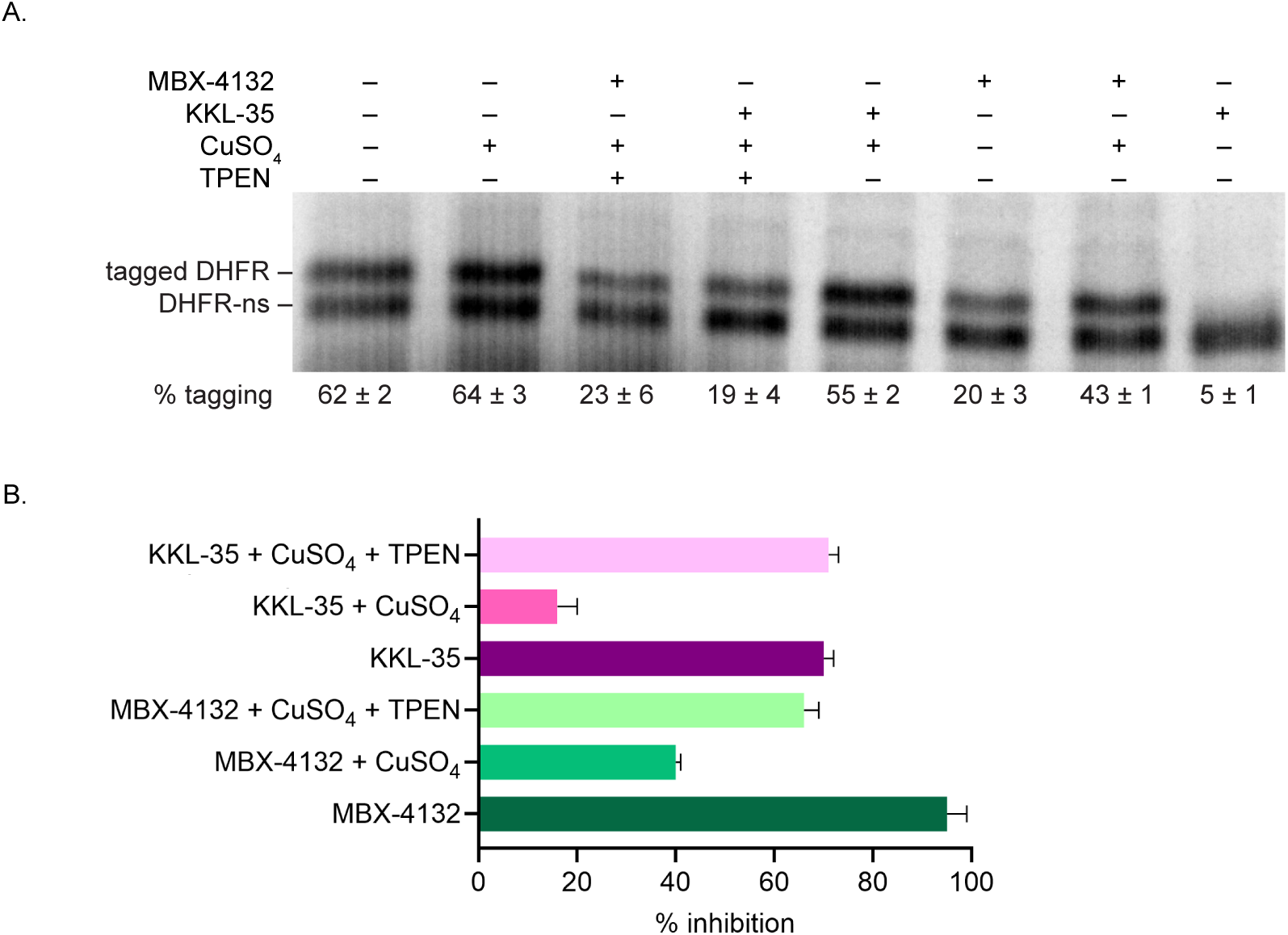
Free copper antagonizes the activity of MBX-4132. A) *in vitro trans*-translation assays as in Figure 2 containing 150 μM CuSO_4_, 150 µM TPEN or 15 μM MBX-4132, as indicated. The average percentage tagging from two independent reactions is shown with the standard deviation. B) Data from gels as in (A) were plotted to show the average from two experiments with error bars indicating the standard deviation.

Although free iron is scarce under infection conditions, if acylaminooxadiazole activity is also inhibited by biologically relevant forms of iron, they would not be clinically useful. However, we found that neither hemin nor transferrin interfered with MBX-4132, MBX-5822A, MBX-6958A, or MBX-4083 activity *in vitro* or during growth inhibition assays (Figure 6, Table 5 & 6). This result is particularly important, as it suggests that the strong antagonism observed with free iron likely will not translate to physiologically relevant conditions. Consequently, these compounds are expected to remain active within the host environment, underscoring their promise as viable candidates for *in vivo* efficacy and future therapeutic development.

**Figure 6.**
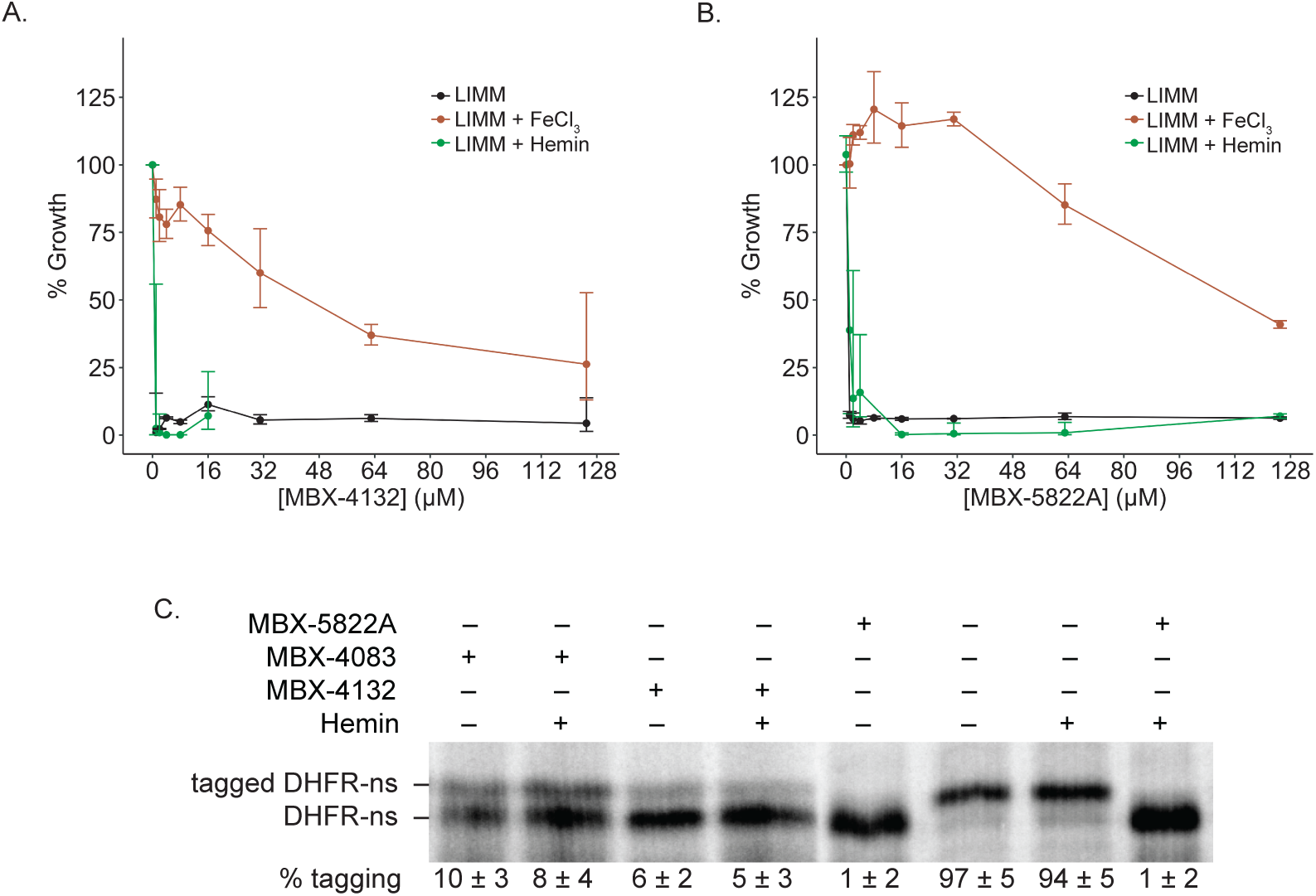
Biologically complexed iron does not decrease activity of acylaminooxadiazoles in cell-based and cell-free assays. A) Growth inhibition by MBX-4132 and MBX-5822A against *M. tuberculosis* H37Rv in LIMM, LIMM supplemented with 40 µM FeCl_3_, and LIMM supplemented with 40 µM hemin. Cells were exposed to acylaminooxadiazoles for 14 days. Data represent geometric means and geometric standard deviations for 3 biological replicates. B) *in vitro trans*-translation assays as in Figure 2 containing 150 μM hemin, 15 µM MBX-5822A or MBX-4132 or MBX-4083, as indicated. The average percentage tagging from two independent reactions is shown with the standard deviation

**Table 5.**
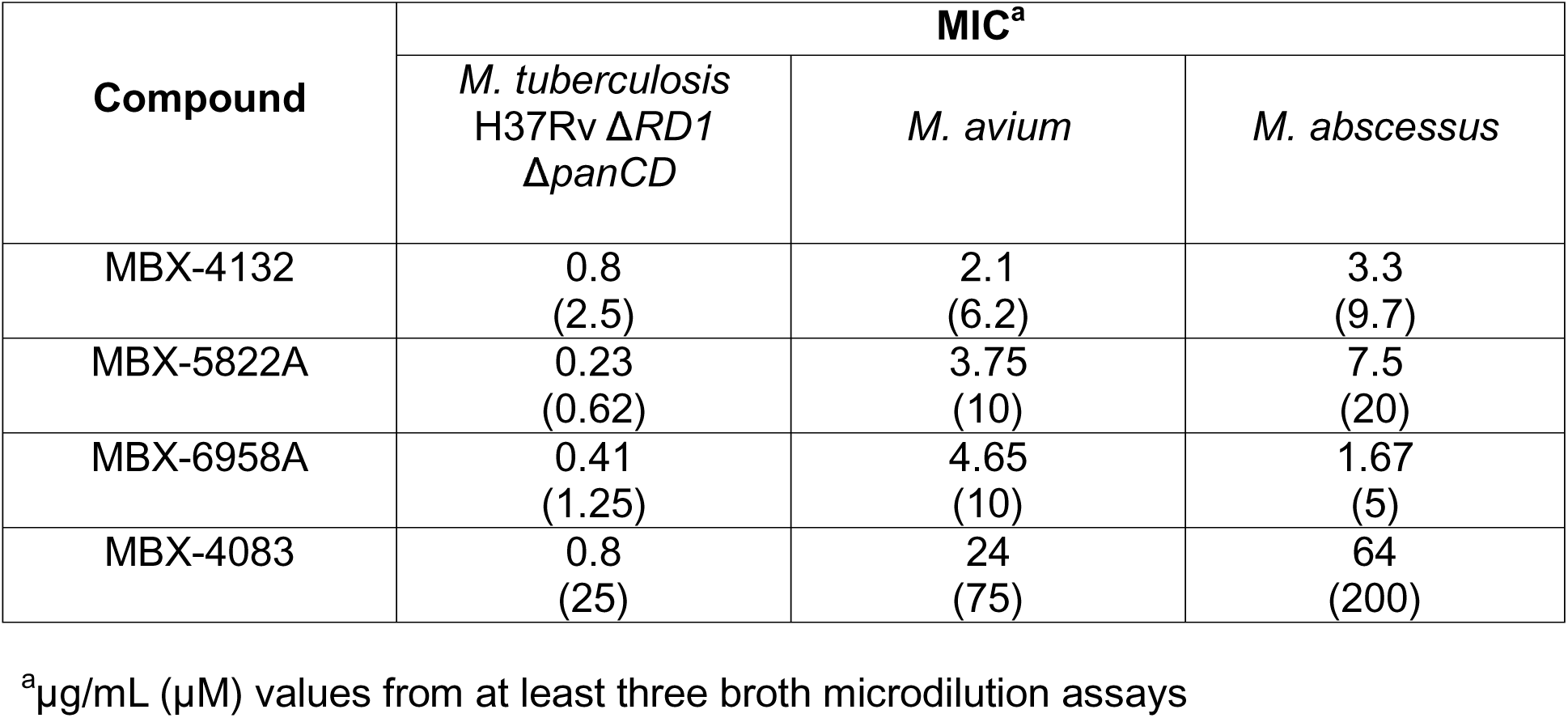
Minimum inhibitory and minimum bactericidal concentrations of *trans-*translation inhibitors for mycobacterial species in LIMM + Hemin.

**Table 6.**
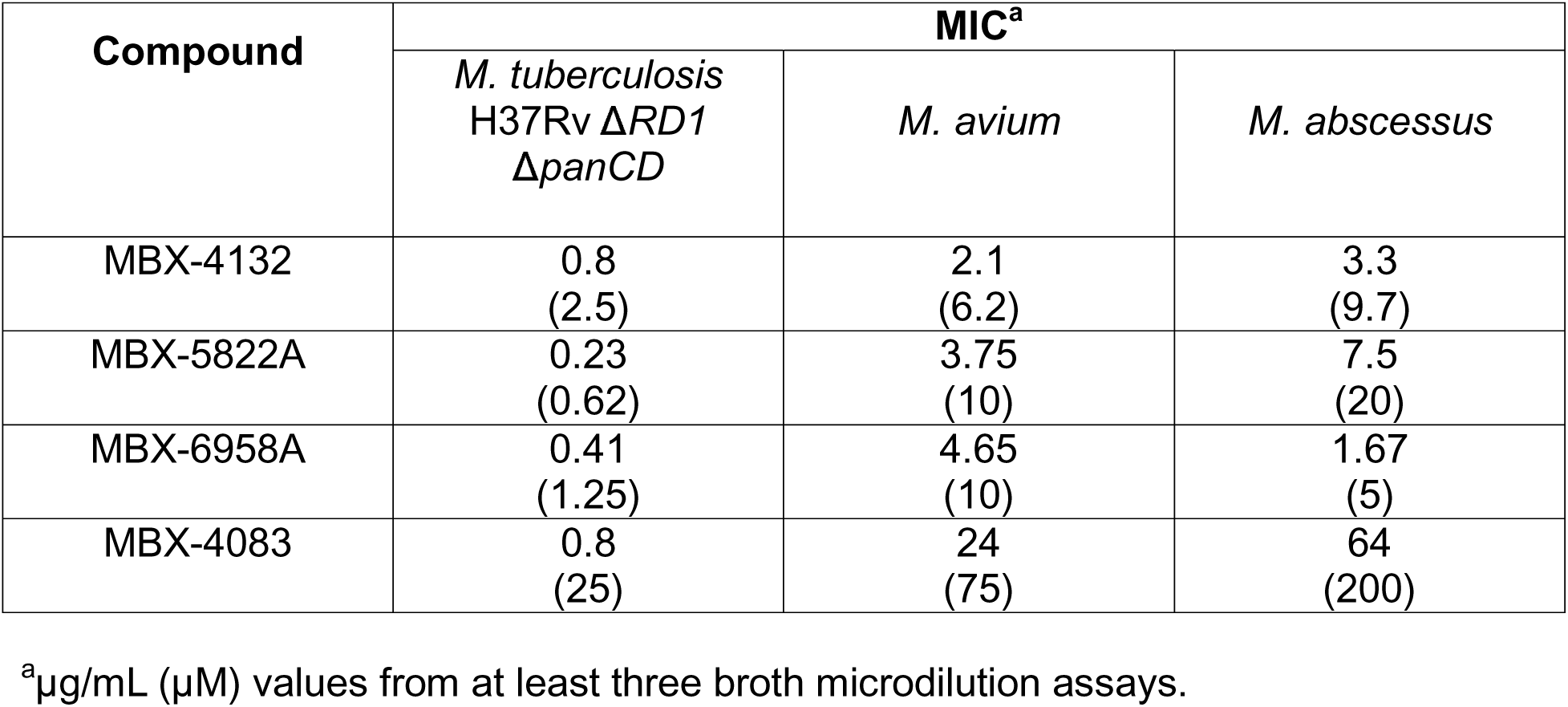
Minimum inhibitory and minimum bactericidal concentrations of *trans-*translation inhibitors for mycobacterial species in LIMM + Holo-transferrin.

### 1,3,4-acylaminooxadiazoles have dual modes of action

Previous transcriptomic data revealed that MBX-4132 causes an iron starvation response in *M. tuberculosis*, suggesting that iron sequestration may enhance biological activity or constitute a separate mechanism of action in addition to inhibition of *trans*-translation [24]. To substantiate the cellular effects of iron starvation caused by MBX-4132, we tested three mutant strains impaired for iron uptake, [34] *mbtF*::Tn (mycobactin biosynthesis), *mmpS5*::Tn (export of deferrated mycobactin), and *irtA*::Tn (import of iron-bound mycobactin and reduction of bound ferric to ferrous) [35]. We observed that all transposon mutants were hypersusceptible to MBX-4132 but not to MBX-4083 (Figure 7A& B).

**Figure 7.**
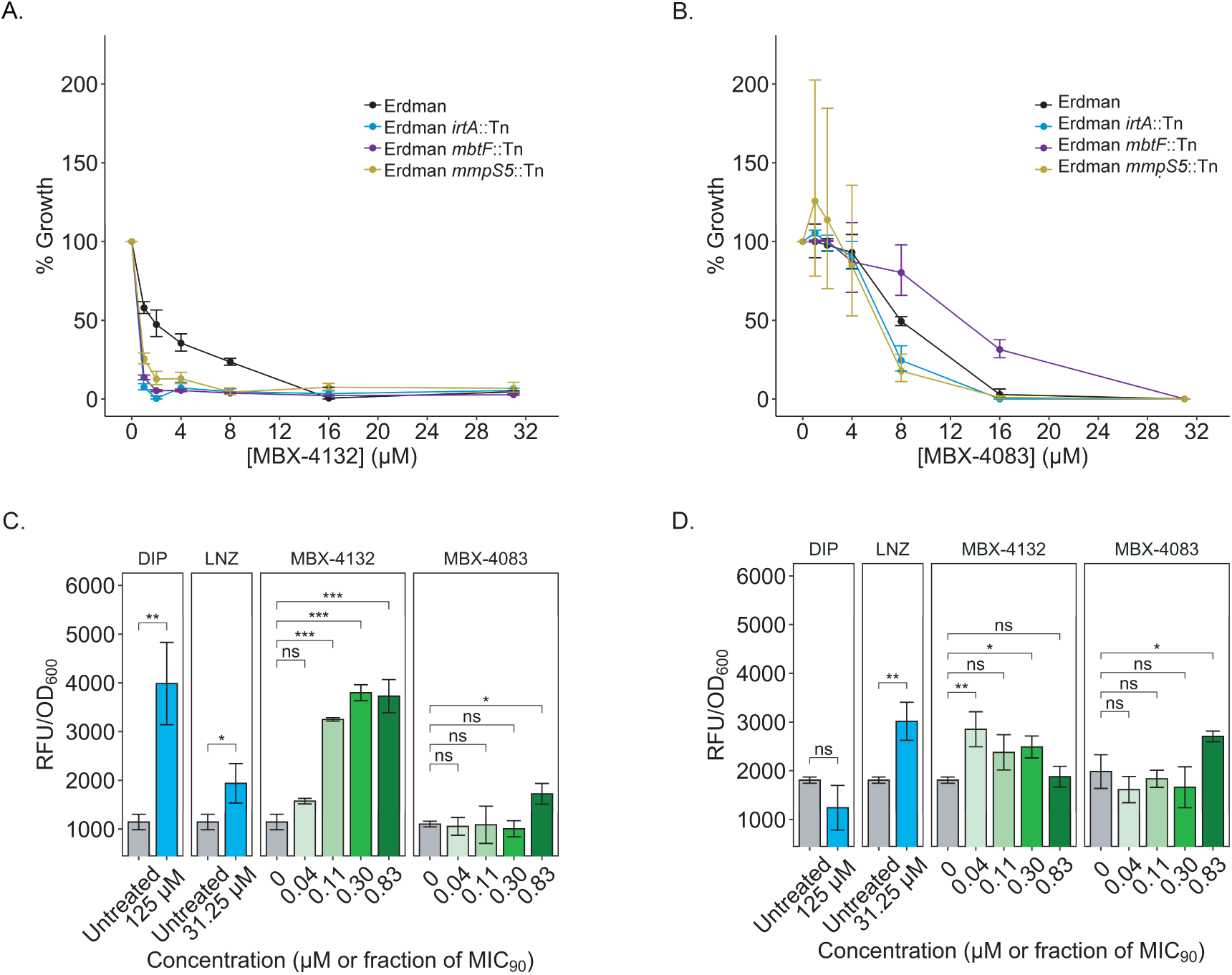
1,3,4-acylaminooxadiazoles disrupt both *trans*-translation and iron homeostasis. A-B) Growth inhibition by (A) MBX-4132 and (B) MBX-4083 against *M. tuberculosis* Erdman and derived Tn mutants in MM following 14 days of treatment. Data represent geometric means and geometric standard deviations for 3 biological replicates. C-D) (C) WhiB7 and (D) MbtH expression level changes as a result of DIP, LNZ, MBX-4132, or MBX-4083 treatment in strains derived from *M. tuberculosis* H37Rv. eGFP signal correlates with expression of the protein of interest and is normalized to bacterial autofluorescence and fluorescence by the media. Cells were treated in HZMM for 48 hours. Molar concentrations of DIP and LNZ were denoted, whereas MBX-4132 and MBX-4083 concentrations were shown as fractions of MIC_90_ in HZMM. Statistical analyses were done by comparing eGFP levels of drug-treated cultures to those of no-drug controls using Dunnett Test. Data represent means and standard deviations of 3 biological replicates. Asterisks indicate statistical significance levels. ns *p* ≥ 0.05, \**p* < 0.05, \*\**p* < 0.01, \*\*\**p* ≤ 0.001.

To determine whether the transcriptional signatures previously observed with MBX-4132 reflected separable cellular responses, we developed fluorescent reporter strains that independently track translation stress and iron starvation. Prior transcriptomic analyses showed that *whiB7* and *mbtH* were strongly induced by MBX-4132, reflecting translational stress and iron starvation, respectively [24, 36, 37]. We therefore constructed *M. tuberculosis* H37Rv strains expressing enhanced green fluorescent protein (eGFP) downstream of *whiB7* and *mbtH* (strains thereby named *whiB7*-*egfp* and *mbtH*-*egfp*, respectively), enabling direct measurement of each pathway in live cells. Reporter responses were assessed using LNZ as a positive control for translation stress and 2,2′-dipyridyl (DIP) as a positive control for iron starvation [37]. MBX-4132 resulted in strong increases in eGFP signals in both *whiB7-egfp* and *mbtH-egfp*, mirroring the dual transcriptional signatures observed previously (Figure 7 C&D) [24]. In contrast, MBX-4083 only led to upregulated eGFP expression in *whiB7-egfp* but not *mbtH-egfp*, even though both compounds were evaluated under identical zinc-rich conditions (Figure 7 C& D). These results indicate that the metal-related transcriptional changes observed with MBX-4132 are not a consequence of *trans-*translation inhibition but represent a distinct activity. Together, these findings demonstrate that MBX-4132 has dual modes of action, inhibition of *trans*-translation and perturbation of iron homeostasis, while MBX-4083 acts only through *trans*-translation inhibition.

## DISCUSSION

This study identifies acylaminooxadiazole analogs with improved activity against *Mycobacterium tuberculosis* and nontuberculous mycobacteria (NTM). Structure–activity relationship analysis showed that changes at both ends of the molecule increased antibacterial potency and *trans*-translation inhibition. Analogs most effective against NTMs, MBX-4370 and MBX-6957, were different from those most potent against *M. tuberculosis.* This separation shows that changes to the acylaminooxadiazoles can shift species specificity. From a therapeutic standpoint, this is useful because treatment for TB and NTM infections is long and creates sustained selective pressure, and any antibiotic used over extended periods will drive resistance. Using different analogs for each pathogen may help limit cross-selection during clinical use and reduce the frequency of resistance emerging.

Because free iron decreased both the antibacterial activity of the 1,3,4-oxadiazoles and their inhibition of *trans-*translation, we infer that these compounds, like MBX-4132 and KKL-35, cause an iron starvation response in addition to targeting *trans-*translation. *M. tuberculosis* mutants defective in siderophore-mediated iron uptake are hypersensitive to these compounds, further supporting this model. Together, these observations suggest that acylaminooxadiazoles influence both *trans-*translation and intracellular metal homeostasis, raising the possibility that they act through more than one mechanism in cells.

However, this free iron-dependent effect is unlikely to limit therapeutic use. In the host, iron is tightly sequestered, and biologically relevant iron sources such as hemin and transferrin did not alter compound activity in culture or *in vitro* assays. Consistent with this, MBX-4132 effectively cleared drug-resistant *Neisseria gonorrhoeae* in mice with a single oral dose, indicating that host iron does not diminish *in vivo* efficacy. If anything, a dual influence on ribosome rescue and metal physiology may be advantageous by requiring bacteria to overcome multiple stresses simultaneously, thereby slowing resistance development.

Our results also show that structural changes enhancing potency do not uniformly affect how the compounds interact with mycobacterial metal physiology. All active 1,3,4-acylaminooxadiazoles inhibit *trans-*translation, but they differ in how they alter iron-regulated pathways. Although the chemical features that separate these activities are not yet defined, the scaffold clearly supports both potent on-target inhibition and a tunable metal-related effect.

## METHODS

### Bacterial Strains, Plasmids, and Growth Conditions

Bacterial strains, plasmids, and primer sequences used in this study are as described previously or listed in Table S4 [24]. *Mycobacterium avium* strain 2285 Smooth, *Mycobacterium abscessus* strain 4530, and *Mycobacterium tuberculosis* H37Rv Δ*RD1* Δ*panCD*, H37Rv, and MTBC clinical isolates were cultured in Middlebrook 7H9 broth (Difco) supplemented with 10% (vol/vol) OADC enrichment (Difco), 0.2% (wt/vol) glycerol, and 0.05% (wt/vol) tyloxapol. Pantothenate was added at a final concentration of 50 mg/L for the auxotrophic strain.

*M. tuberculosis* H37Rv, Erdman, H37Ra, and derived strains were cultured in Mtb Minimal Medium (MM), LIMM, LIMM containing FeCl_3_ or hemin, HZMM [24], or HZMM containing CuSO_4_, as consistent with downstream MIC assays.

*Mycobacterium avium*, *Mycobacterium abscessus*, and *M. tuberculosis* H37Rv ΔRD1 Δ*panCD* were plated on Middlebrook 7H10 agar (Difco) containing 10% (vol/vol) OADC enrichment, 0.2% (wt/vol) glycerol. Pantothenate (50 mg/L) was added wherever required. *Escherichia coli* strains were grown in Lysogeny Broth (LB; Difco). *E. coli* DH5α was used for plasmid propagation, while *E. coli* BL21(DE3) was used for overexpression and purification of *M. tuberculosis* translation factors and grown in LB supplemented with 50 μg/mL kanamycin.

### MIC, MBC, and Synergy Assays

MIC values were determined by broth microdilution in 96-well plates according to CLSI guidelines for non-tuberculosis mycobacteria (NTM) and attenuated *M. tuberculosis* [38]. Plates were incubated at 37 °C for 3 days for *M. avium* and *M. abscessus,* and 1 week for *M. tuberculosis.* The MIC was recorded as the lowest compound concentration that prevented visible growth. All assays were performed in at least three independent biological replicates. To determine MBC values, 5 µL from wells corresponding to the MIC, 2× MIC, and 4× MIC were spotted onto 7H10 agar or LB agar and incubated under the same conditions. A compound was considered bactericidal if it produced ≥ 99 % reduction in CFU at 2× MIC and no colonies at 4× MIC. The MBC was defined as the lowest concentration yielding no CFU.

MIC values in virulent *M. tuberculosis* strains were determined using the modified MTT [3-(4,5-dimethylthiazol-2-yl)-2,5-diphenyltetrazolium bromide] assay as previously illustrated by our group [24]. Cells were subjected to 7 days of acylaminooxadiazole treatment in Middlebrook 7H9 or 7H9 containing 100 µM ZnSO_4_ and 14 days of treatment in MM and other MM-based media unless otherwise noted. Susceptibility of CRISPR (Clustered Regularly Interspaced Short Palindromic Repeats) interference (CRISPRi) mutants, which had been constructed as part of our previous study, to acylaminooxadiazoles was assessed as previously described [24].

Synergy between MBX compounds and clinically relevant agents was evaluated in LIMM using a checkerboard microdilution format. Two-fold serial dilutions of each compound were arrayed along the plate axes, and plates were incubated at 37 °C for 3 days for *M. avium* and *M. abscessus,* and 1 week for *M. tuberculosis.* The fractional inhibitory concentration (FIC) for each compound was calculated as the MIC of the compound in combination divided by its MIC alone. The fractional inhibitory concentration index (FICI = FIC_A + FIC_B) was used to classify interactions as synergistic (FICI ≤ 0.5), additive (0.5 < FICI ≤ 1.0), indifferent (1 < FICI ≤ 4.0), or antagonistic (FICI > 4.0)

### *In vitro* translation and *trans*-translation assays

*M. tuberculosis trans*-translation assays were performed using purified *M. tuberculosis* components based on previous protocols [24, 26]. Energy solution and *M. tuberculosis* protein solution were prepared as described previously [24, 26]. *M. tuberculosis* tmRNA was transcribed *in vitro* using primers MTB ssrA F and MTB ssrA R, and 6×His-tagged *M. tuberculosis* SmpB and EF-Tu were purified from *E. coli* BL21(DE3) carrying pET28aTBsmpB-His₆ and pET28aTBuf-His₆, respectively, as previously reported [24, 26]. *M. tuberculosis* ribosomes were isolated from H37Rv ΔRD1 Δ*panCD* as described before [24, 26].

For translation assays, reactions were assembled with the energy solution (2 µL), *M. tuberculosis* protein solution (1 µL), *M. tuberculosis* EF-Tu (10 µM), *M. tuberculosis* ribosomes (1.28 µM), a *dhfr-stop* DNA template (9 ng/µL), and [³ S]-methionine (0.42 µCi/µL). Reactions were incubated at 37 °C for 2 h, precipitated with acetone, resolved by 15 % SDS-PAGE, and visualized by phosphor imaging (GE Healthcare, Chicago, IL, USA). Wherever indicated, 0.25 µL of MBX-5822A, MBX-6958A, MBX-4083, chloramphenicol, or dimethyl sulfoxide (DMSO) was added. Relative translation activity was determined relative to the DMSO control.

For *trans*-translation assays, reactions were assembled as described previously [24, 26]. The *dhfr-ns* template lacking an in-frame stop codon was prepared by PCR and added to a final concentration of 9 ng/µL. Purified *M. tuberculosis* tmRNA and SmpB were included at a final concentration of 2.75 µM each. Reactions were incubated at 37 °C for 2 h, precipitated with acetone, separated by 15 % SDS-PAGE, and analyzed by phosphor imaging.

To evaluate the effect of iron or copper, Fe₂(SO₄)₃ or CuSO_4_ water was pre-incubated with MBX-5822A, MBX-6958A, MBX-4083, or DMSO for 10 min at room temperature before addition to the reaction mixture. Unless otherwise stated, final concentrations were Fe₂(SO₄)₃ (150 µM), CuSO_4_ (150 µM), MBX-5822A (15 µM), MBX-6958A (15 µM), and MBX-4083 (30 µM). Tagging efficiency was defined as the percentage of total DHFR that acquired the tmRNA-encoded peptide tag and averaged from three independent experiments. Dose-response curves were generated from at least three replicates, and IC₅₀ values were calculated using nonlinear regression fits in GraphPad Prism.

### *M. tuberculosis* mutant construction

*M. tuberculosis* H37Rv *whiB7*-*egfp* and *mbtH*-*egfp* strains were created using <u>o</u>ligonucleotide-mediated recombineering followed by <u>B</u>xb1 integrase targeting (ORBIT) [39] with the targeting oligonucleotides shown in Table S4 and co-transformed with pUMN105 (Addgene #233024) using ORBIT protocols previously described [24]. Successful integration of pUMN105 was confirmed using PCR.

### Fluorescence assays

*M. tuberculosis* H37Rv *whiB7-egfp* and *mbtH-egfp* were cultured in HZMM until log-phase at 37 °C with shaking. Cells were diluted to an OD_600_ of 0.01 in the same medium and exposed to 31.25 µM LNZ, 125 µM DIP, MBX-4132 or MBX-4083, or equal-volume DMSO for 2 days. These cultures were incubated at 37 °C without shaking in 96-well glass-bottom microplates (Greiner Bio-One, SensoPlate). Fluorescence was measured using a microplate reader (BioTek Synergy H1) and normalized to OD_600_. Dunnett’s Test was used for statistical analyses, comparing drug-treated samples with respective DMSO controls, with *p*-values < 0.05 considered statistically significant.

## ACKNOWLEDGEMENTS

We thank William R. Jacobs, Jr. of the Albert Einstein College of Medicine for providing *M. tuberculosis* strains H37Rv and H37Rv Δ*panCD*Δ*RD1*. We are grateful to the NIH NIAID for financial support of this project through grant AI158706. We are also grateful to the Biosafety Level 3 Program for providing essential institutional support for this project.

