## Supporting Information for "*trans*-Translation inhibitors that kill *M. tuberculosis* and pathogenic Non-tuberculous Mycobacteria have a dual mechanism of action"

**AUTHOR AFFILIATIONS AND FOOTNOTES:**

[1] Department of Molecular Biosciences, The University of Texas at Austin, Austin, TX 78712 USA.

[2] Department of Microbiology and Immunology, University of Minnesota Medical School, Minneapolis, MN 55455 USA.

[3] Microbiotix, Inc., Worcester, MA 01605 USA

[4] LaMontagne Center for Infectious Diseases, University of Texas at Austin, Austin, TX

[‡] These authors contributed equally

### SUPPORTING INFORMATION

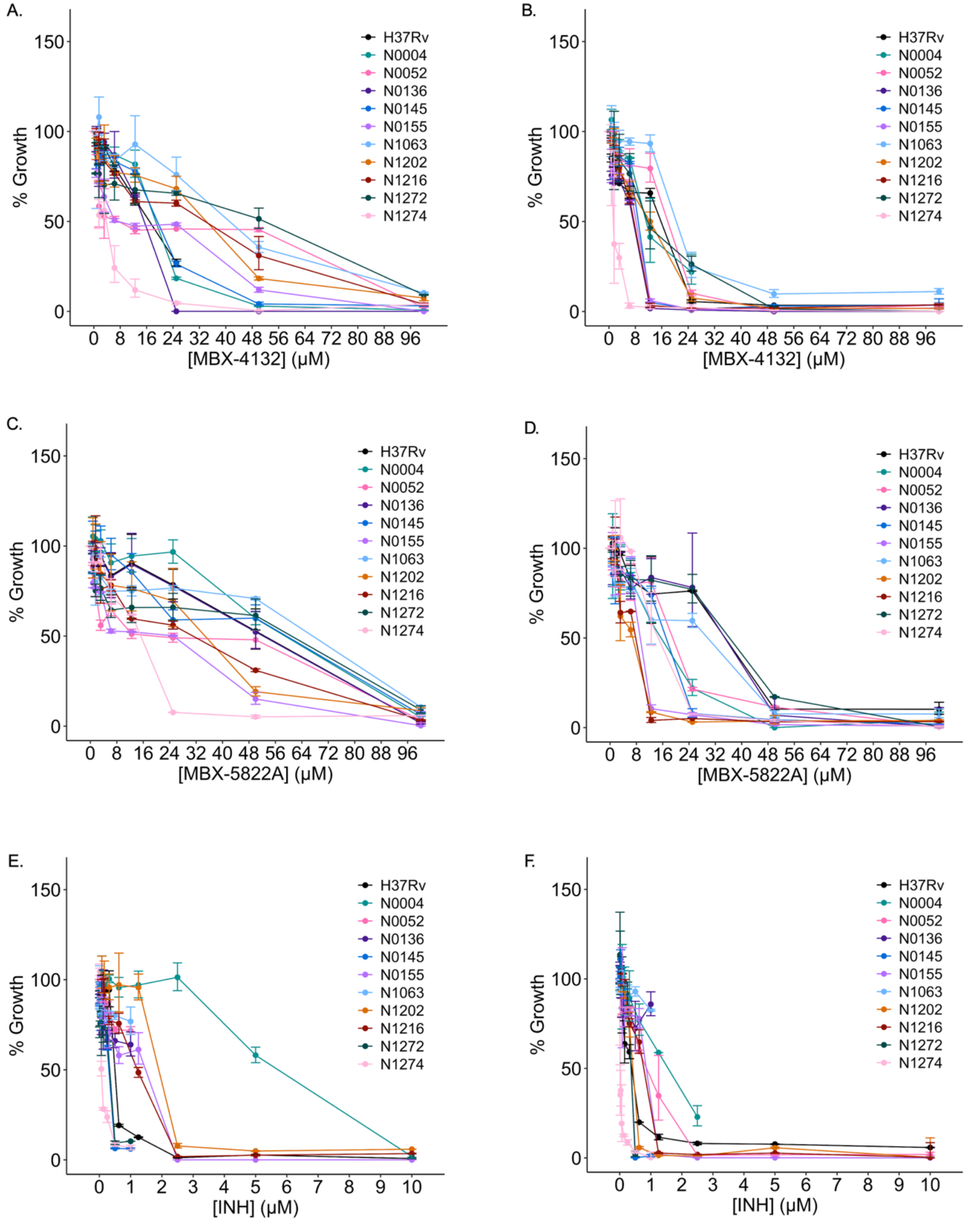


**Figure S1. Acylaminooxadiazole inhibition against MTBC clinical isolates.** Cells were treated with (A-B) MBX-4132, (C-D) MBX-5822A, or (E-F) INH for 7 days in (A,C,E) Middlebrook 7H9 or (B,D,F) 7H9 supplemented with 100 µM $ZnSO_4$. Data represent geometric means and geometric standard deviations for 2 biological replicates.

### SUPPLEMENTAL TABLES

**Table S1. Minimum inhibitory and minimum bactericidal concentrations of *trans*-translation inhibitors for mycobacterial species in 7H9 medium**

| Compound | *M. tuberculosis* H37Rv Δ*RD1* Δ*panCD* | | *M. avium* | | *M. abscessus* | |
|---|---|---|---|---|---|---|
| | **MIC[a]** | **MBC[b]** | **MIC** | **MBC** | **MIC** | **MBC** |
| KKL-35 | >40 (>125) | >40 (>125) | >40 (>125) | >40 (>125) | >40 (>125) | >40 (>125) |
| MBX-4132 | >40 (>125) | >40 (>125) | >40 (>125) | >40 (>125) | >40 (>125) | >40 (>125) |
| MBX-5822A | 40 (125) | 40 (125) | 40 (125) | 40 (125) | >40 (>125) | >40 (>125) |
| MBX-6958A | 40 (125) | 40 (125) | 40 (125) | >40 (>125) | >40 (>125) | >40 (>125) |
| MBX-6957 | >40 (>125) | >40 (>125) | 40 (125) | 40 (125) | >40 (>125) | >40 (>125) |
| MBX-4370 | >40 (>125) | >40 (>125) | 40 (125) | 40 (125) | 40 (125) | 40 (125) |
| MBX-4083 | 0.8 (25) | 0.8 (25) | 24 (75) | 24 (75) | 64 (200) | 64 (200) |
| KKL-38989 | 32 (100) | 32 (100) | >32 (>100) | >32 (>100) | >32 (>100) | >32 (>100) |
| KKL-39893 | 16 (50) | 16 (50) | >32 (>100) | >32 (>100) | >32 (>100) | >32 (>100) |
| KKL-39894 | 32 (100) | 32 (100) | >32 (>100) | >32 (>100) | >32 (>100) | >32 (>100) |
| KKL-39895 | 32 (100) | 32 (100) | >32 (>100) | >32 (>100) | >32 (>100) | >32 (>100) |
| KKL-39896 | 32 (100) | 32 (100) | >32 (>100) | >32 (>100) | >32 (>100) | >32 (>100) |
| Rifampicin | 0.12 (0.14) | 0.12 (0.14) | 0.5 (0.6) | 1 (1.2) | 128 (155) | 256 (311) |
| Azithromycin | 4 (5.3) | ND[c] | 32 (42.4) | ND | 8 (10.6) | ND |

[a]µg/mL (µM) values from at least three broth microdilution assays.

[b]µg/mL (µM) values from at least three plating assays.

[c]Not determined.

**Table S2. Effect of $Cu^{2+}$ on antibacterial activity of acylaminooxadiazoles in LIMM**

| Compound | *M. tuberculosis* H37Rv Δ*RD1* Δ*panCD* | | *M. avium* | | *M. abscessus* | |
|---|---|---|---|---|---|---|
| | **MIC[a]** | **MBC[b]** | **MIC** | **MBC** | **MIC** | **MBC** |
| KKL-35 | >40 (>125) | >40 (>125) | >40 (>125) | >40 (>125) | >40 (>125) | >40 (>125) |
| MBX-4132 | >40 (>125) | >40 (>125) | >40 (>125) | >40 (>125) | >40 (>125) | >40 (>125) |
| Rifampicin | 0.12 (0.14) | 0.12 (0.14) | 0.5 (0.6) | 1 (1.2) | 128 (155) | 256 (311) |
| Azithromycin | 4 (5.3) | ND[c] | 32 (42.4) | ND | 8 (10.6) | ND |

[a]μg/mL (μM) values from at least three broth microdilution assays.

[b]μg/mL (μM) values from at least three plating assays.

[c] Not determined.

**Table S3. Effect of $Cu^{2+}$ on antibacterial activity of acylaminooxadiazole in HZMM**

| Compound | MIC in MM[a] | MIC in HZMM[a] | MIC in HZMM + 1 μM $CuSO_4$[a] |
|---|---|---|---|
| MBX-4132[b] | >80 (>250) | 1.3 (4) | >80 (>250) |
| MBX-5822A | >80 (>250) | <0.3 (<1) | >80 (>250) |
| MBX-6958A | >80 (>250) | <0.3 (<1) | >8 (>25) |
| MBX-4083 | 1.0 (31) | 1.0 (31) | 0.5 (16) |

[a]μg/mL (μM) values from at least three broth microdilution assays against *M. tuberculosis* H37Rv.

[b]MBX-4132 MICs in MM and HZMM were described previously (25).

**Table S4. Bacterial strains and plasmids**

| Strain | Description | Source or reference |
|---|---|---|
| *M. tuberculosis* N0004 | Clinical isolate, lineage 3 from India | Gift from Berney Lab (Borrell *et al*., 2019) |
| *M. tuberculosis* N0052 | Clinical isolate, lineage 2.2.2 from China | Gift from Berney Lab (Borrell *et al*., 2019) |
| *M. tuberculosis* N0136 | Clinical isolate, lineage 4.3.3 from USA | Gift from Berney Lab (Borrell *et al*., 2019) |
| *M. tuberculosis* N0145 | Clinical isolate, lineage 2.2.1.1 from China | Gift from Berney Lab (Borrell *et al*., 2019) |
| *M. tuberculosis* N0155 | Clinical isolate, lineage 2.2.1 from China | Gift from Berney Lab (Borrell *et al*., 2019) |
| *M. africanum* N1063 | Clinical isolate, lineage 5, requires 40 mM pyruvate | Gift from Berney Lab (Guerrero-Bustamante *et al*., 2021) |
| *M. africanum* N1202 | Clinical isolate, lineage 6 from Ghana, requires 40 mM pyruvate | Gift from Berney Lab (Borrell *et al*., 2019) |
| *M. tuberculosis* N1216 | Clinical isolate, lineage 4.6.2.2 from Ghana | Gift from Berney Lab (Borrell *et al*., 2019) |
| *M. africanum* N1272 | Clinical isolate, lineage 5 from Ghana, requires 40 mM pyruvate | Gift from Berney Lab (Borrell *et al*., 2019) |
| *M. tuberculosis* N1274 | Clinical isolate, lineage 3 from Afghanistan, streptomycin resistant | Gift from Berney Lab (Borrell *et al*., 2019) |
| *M. tuberculosis* Erdman | WT strain; virulent | Gift from Tischler Lab (Block *et al*., 2023) |
| *M. tuberculosis* Erdman *mbtF*::Tn | Erdman derivative; contains transposon insertion in *mbtF* | Gift from Tischler Lab (Block *et al*., 2023) |
| *M. tuberculosis* Erdman *mmpS5*::Tn | Erdman derivative; contains transposon insertion in *mmpS5* | Gift from Tischler Lab (Block *et al*., 2023) |

| | | |
|---|---|---|
| *M. tuberculosis* Erdman *irtA*::Tn | Erdman derivative; contains transposon insertion in *irtA* | Gift from Tischler Lab (Block *et al*., 2023) |
| *M. tuberculosis* H37Rv *whiB7-egfp* | H37Rv derivative; pUMN105 integrated downstream of *whiB7* | This study |
| *M. tuberculosis* H37Rv *mbtH-egfp* | H37Rv derivative; pUMN105 integrated downstream of *mbtH* | This study |
| *M. tuberculosis* H37Ra | WT strain; avirulent | Gift from Jacobs Lab |
| *M. tuberculosis* H37Ra pKM461 | H37Ra derivative; expresses RecT annealase and Bxb1 integrase; tetracycline-inducible | Previously constructed in Baughn Lab |
| *E. coli* HB101 | For propagation of plasmids | (Glover, 1985) |
| *E. coli* HB101 pUMN105 | Contains plasmid pUMN105 | This study |

| **Plasmid** | **Description** | **Source or reference** |
|---|---|---|
| pUMN105 | pKM468 derivative; expresses enhanced green fluorescent protein (eGFP); $hyg^R$ | This study |

| **Oligonucleotide** | **Sequence 5' to 3'** | **Source or reference** |
|---|---|---|
| *whiB7-egfp* ORBIT targeting oligo | CGGGTTTCGCCGAGCCCGACGCGATCGTCGTCTGAGCCGGCTCGCGCCGGCGGGCGCACCATCGCGGGGGTTTGTACCGTACACCACTGAGACCGCGGTGGTTGACCAGACAAACCCATCCTCCTTCCTCCTTCTATGCAACAGCATCCTTGCGCGGACGTCCGCGCGGACGCTTGTGACTCACGATCGAGCCTTGGTCG | This study |
| *mbtH-egfp* ORBIT targeting oligo | GATTGTGCGTTTGGTTCAAACTACCGGTGCACGCGCCGGGCAAGTCTGTGTCGTAGCTGTGAGCGAGCGGGGTTTGTACCGTACACCACTGAGACCGCGGTGGTTGACCAGACAAACCCATCCTCCTTCCTCCTTTCAGTCCTCGACCATGGCGTCACGCAGGCTCTTCGGCCGCAGATCGGTCCAGTTCTTTTCCACGT | This study |
